# Reactive Oxygen Species Generation Drives Iron Accumulation by Rotenone-Mediated Inhibition of Mitochondrial Complex I in Dopaminergic Neurons

**DOI:** 10.64898/2026.09.04.749533

**Authors:** Alaa S. Abdelrazeq Hassan, Saniya Pungliya, Matias A. Murillo, Karlee H. Parry, Amy R. Apfelbaum, Anthony S. Grillo

## Abstract

Iron is an essential element that plays a critical role in mitochondrial bioenergetics, yet its excess is cytotoxic and contributes to the development of neurodegenerative diseases such as Parkinson’s disease (PD). Impaired Complex I (CI) activity, which can be caused by exposure to the environmental toxin rotenone, is implicated in PD. The mechanistic underpinnings connecting iron dyshomeostasis to cytotoxicity remain unclear, which creates challenges towards preventing neuronal cell death in PD. We hypothesized that CI inhibition is sufficient to disrupt cellular iron homeostasis via reactive oxygen species (ROS) generation. Using SH-SY5Y cells differentiated into dopaminergic neurons, we show that rotenone-induced CI inhibition increases ROS, promotes oxidative stress and cytotoxicity, and drives a redistribution of labile iron. Specifically, mitochondrial and total cellular iron levels increase, while cytosolic labile iron is reduced. Antioxidant treatment blocks both ROS production and iron accumulation, suggesting that ROS is critical for iron maldistribution in our system. Conversely, iron chelation suppresses ROS propagation, suggesting a positive feedback loop in which iron further amplifies oxidative stress. Our data are consistent with a greater sensitivity of mitochondrial [4Fe-4S]-containing proteins relative to the [2Fe-2S] proteins examined, potentially contributing to mitochondrial iron retention. Together, these findings establish a mechanistic link between mitochondrial dysfunction and iron dyshomeostasis and offer insights into how environmental CI inhibitors contribute to the pathogenesis of PD. More broadly, this work may have relevance to sporadic PD and other genetic or age-related disorders associated with iron accumulation and mitochondrial diseases such as Leigh Syndrome.

## Introduction

Mitochondrial Complex I deactivation is increasingly recognized as pathogenic in Parkinson’s disease (PD) and other age-related neurodegenerative diseases (1–3). Further, mutations in Complex I subunits or other proteins in the electron transport chain (ETC) cause pediatric mitochondrial diseases such as Leigh Syndrome that display parkinsonism-like features (2, 4–6). Exposures to mitochondrial poisons, including the pesticide rotenone, are thought to be responsible for many environmentally induced cases of PD (7–10). However, the mechanistic underpinnings of how rotenone effects biomolecular pathways to cause cellular toxicity remains unclear. Discovering the pathobiology of iron-mediated toxicity thus presents an opportunity to better understand environmental risk factors that cause PD. Notably, our prior work suggests Complex I inhibition is associated with iron accumulation, a common hallmark of PD (1, 6).

Iron is a crucial element in humans that plays a key role in oxygen transport, the immune response, and metabolism, among other functions (11). Iron exists in biological systems in multiple forms, with a large prevalence of forming iron-sulfur clusters (ISCs). ISCs are essential for electron transport in the ETC, and ISC-containing proteins are vital in many other metabolic and biochemical processes (12–15). Despite its primary and essential role in cells, iron progressively accumulates in the body with age and in many neurodegenerative diseases (16, 17). This excess iron promotes the generation of reactive oxygen species (ROS), which have been implicated in virtually every age-related disease and in normative aging by inducing oxidative damage. Because of this, excess iron is established as a hallmark of some neurodegenerative diseases, including PD (1). In PD patients, elevated iron levels in the substantia nigra of the brain correlate well with disease severity (18–20). Because of the inherent and damaging reactivity of excess iron through the Fenton reaction, iron is tightly regulated at the systemic and cellular levels.

Iron homeostasis is maintained at the transcriptional, translational, and post-translational level through independent mechanisms (21). Circulating hormones such as hepcidin influence body iron absorption and cellular iron excretion by promoting the degradation of the iron export protein ferroportin (FPN1) (11, 22). On the cellular level, the iron-regulatory proteins IRP1 and IRP2 sense cytosolic labile iron to dissociate from iron-responsive elements on the 5’- or 3’-untranslated regions of the iron-related protein transcripts transferrin receptor 1 (*TFR1*), divalent metal transporter 1 (*DMT1*), ferritin subunits (*FTL1/FTH1*), and *FPN1,* amongst others. Upon iron recognition and binding, IRP1 functionally switches to act as a cytosolic aconitase. IRP2 is ubiquitinated and degraded after recognition by the ISC protein FBXL5. Cellular iron balance is further regulated by transcription factor activation such as the hypoxia inducible factors (HIF1α and HIF2α) or NRF2 in response to iron status and oxidative stress (23–26). Many environmental stressors can cause this oxidative stress, generate ROS, and induce the improper regulation or iron. However, because of the complex and interdependent regulation of iron homeostasis, the subcellular sequence of interactions leading to iron misregulation are unclear and often overlooked (27).

ROS are byproducts of aerobic metabolism and normally play essential roles in cell signaling, growth, and metabolic adaptation (28). However, excess, unchecked ROS generation caused by environmental toxins such as rotenone or genetic factors is often cytotoxic. Recent literature suggests that ROS production, mitochondrial dysfunction, and iron dyshomeostasis are connected contributors to PD pathogenesis rather than independent disease features (2, 6, 17). Genetic disruption of mitochondrial Complex I (via *Ndufs2* deletion) in dopaminergic neurons is sufficient to produce levodopa-responsive progressive parkinsonism-like pathology in mice (2), while human quantitative susceptibility mapping and other studies show iron accumulation and/or deposition with PD progression (19, 20). This supports a close relationship between mitochondrial dysfunction, dopaminergic toxicity, and iron homeostasis. Mechanistically, iron can amplify oxidative stress by inducing lipid peroxidation and cell death (29–31). In rotenone-based PD models, the relationship between ROS and iron appears especially relevant, as rotenone-induced dopaminergic neurotoxicity is associated with iron imbalances, lipid peroxidation, mitochondrial morphological alterations, and NCOA4-dependent ferritinophagy (22, 32, 33). Together, these studies support the idea that perturbed iron mobilization may occur downstream of mitochondrial dysfunction and contribute to the propagation of oxidative damage.

More broadly, as oxidative damage is implicated in virtually every neurodegenerative disease, it is essential to better understand the molecular sequence of events by which excess ROS production induces oxidative damage and cellular toxicity. As excess ROS generation, disturbed iron metabolism, and mitochondrial Complex I dysfunction are widely considered hallmarks of PD (1), we asked whether they are tightly connected. Specifically, we hypothesized that Complex I deactivation by the environmental PD risk factor rotenone perturbs iron homeostasis through the generation of reactive oxygen species. Herein, we use differentiated dopaminergic neurons to demonstrate that rotenone induces iron accumulation, highlighted by mitochondrial iron retention with a cytosolic iron deficiency. Further, we show that ROS generation by rotenone is essential for iron misregulation, as antioxidant treatment prevents iron imbalances. Collectively, this work leads to a better understanding of the molecular sequence events by which rotenone causes neuron cytotoxicity using *in vitro* PD models.

## Results

### Rotenone induces iron accumulation in neurons

PD causes dopaminergic neuron cell death and is associated with the accumulation of iron in these cells. To mechanistically study the behavior of iron in an *in vitro* PD model, we first chemically differentiated SH-SY5Y neuroblastoma cells into dopaminergic neurons using retinoic acid and TPA phorbol ester using established protocols (34). We confirmed high enrichment of dopaminergic neurons after differentiation by probing cells positively stained for the dopaminergic marker tyrosine hydroxylase (TH) using immunofluorescence labeling (Fig. S1). We also observed increased expression of MAP2 (Fig. S1A), consistent with neuron differentiation. Following literature precedent, we next treated cells with the selective and potent Complex I inhibitor rotenone to represent an effective PD cell model (8–10). Iron is reported to accumulate in the substantia nigra of PD patient brains (18, 19). We thus asked whether rotenone treatment induces the accumulation of iron in our system. Since the non-heme iron represents most of the iron content in neurons, we quantified the non-heme iron levels in vehicle- and rotenone-treated cells using a ferrozine colorimetric assay (35). We observed a significant increase in non-heme iron in rotenone-treated cells relative to control (Fig. 1A). We used inductively coupled plasma-mass spectrometry (ICP-MS) to quantify total iron levels in vehicle- and rotenone-treated cells. Consistent with our non-heme iron analysis, we observed that rotenone treatment also increased total cellular iron status (Fig. 1B).

**Figure 1.**
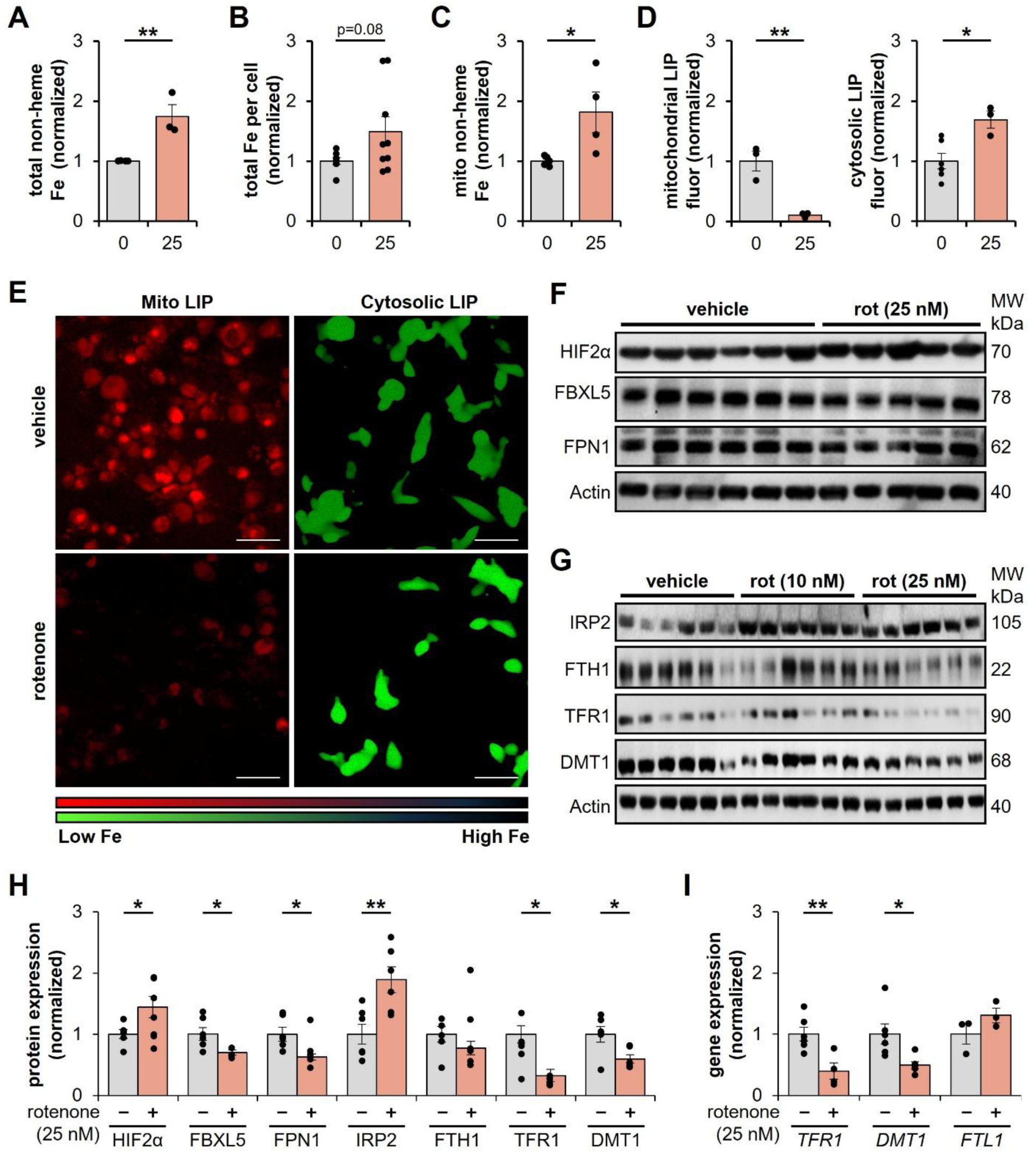
Rotenone causes iron maldistribution in neurons. **(A)** Total non-heme iron levels via ferrozine colorimetric assay in differentiated SH-SY5Y cells treated with rotenone (25 nM) or DMSO (0 nM) for 24 hrs. **(B)** Total iron levels via ICP-MS in differentiated SH-SY5Y cells treated with rotenone (25 nM) or DMSO (0 nM) for 24 hrs. **(C)** Non-heme iron levels via ferrozine colorimetric assay in isolated mitochondrial pellets from differentiated SH-SY5Y cells treated with rotenone (25 nM) or DMSO (0 nM) for 24 hrs. **(D**, **E)** Assessment of changes in labile iron pools in the mitochondria and cytosol in differentiated SH-SY5Y cells treated with rotenone (25 nM) or DMSO vehicle (0 nM) using the iron-responsive fluorescent dyes RPA (mitochondria, red) and Calcien Green-AM (cytosol, green). Scale bars = 50 µm. **(F**, **G)** Representative images of western blot of proteins involved in iron homeostasis and iron regulation in differentiated SH-SY5Y neurons treated with DMSO vehicle (0 nM) or rotenone (10 or 25 nM) for 24 hrs. **(H)** Densitometry of iron-related protein expression relative to actin and normalized to control levels for each individual protein in differentiated SH-SY5Y neurons treated with DMSO (0 nM) or rotenone (25 nM) for 24 hrs. **(I)** Gene expression levels of IRE-responsive genes by qPCR relative to *GAPDH* expression and normalized to controls for each gene analyzed in differentiated SH-SY5Y neurons treated with DMSO (0 nM) or rotenone (25 nM) for 24 hrs. *p < 0.05, **p < 0.01

Our total cellular iron analysis was unable to provide information on the intracellular distribution of iron inside of organelles such as the mitochondria. We thus asked whether the increase in iron is attributed to global cellular iron retention, or if it is driven by the maldistribution of intracellular iron. In other words, we simply asked whether rotenone treatment affects cellular iron distribution. Mitochondria contain a significant portion of total cellular iron due to the demanding role iron plays in mitochondrial metabolism, heme synthesis, and ISC biogenesis (13, 14, 36). We thus isolated mitochondria using differential centrifugation in vehicle- and rotenone-treated SH-SY5Y cells to ask whether rotenone promotes mitochondrial iron retention. Mitochondrial enriched fractions were confirmed by immunoblot targeting the mitochondrial-localized protein TOMM20 (Fig. S2). We then analyzed mitochondrial non-heme iron levels in vehicle- or rotenone-treated cells after isolating mitochondria with a ferrozine assay. We observed mitochondrial non-heme iron-accumulation in rotenone-treated cells (Fig. 1C), similar to our results showing total cellular iron accumulation.

### Rotenone causes the uneven distribution of mitochondrial and cytosolic labile iron

As most of the cellular iron is either protein-bound or in a more reactive and dynamic form referred to as labile iron that can generate ROS through Fenton chemistry in excess (30, 37), we next evaluated whether the mitochondrial iron retention increases mitochondrial labile iron. To achieve this, we used the iron-responsive fluorescent dye RPA in vehicle- and rotenone-treated neurons. We observed a quenching of fluorescence upon rotenone treatment (Fig. 1D, 1E and Fig. S3), consistent with increased labile iron in mitochondria. We next used Calcein Green-AM, whose fluorescence quenches upon binding iron in the cytosol. In contrast to that observed with mitochondrial LIP, rotenone decreased cytosolic labile iron as indicated by an increase in Calcein Green fluorescence (Fig. 1D, 1E and Fig. S3). This surprising result of mitochondrial iron retention and cytosolic iron deficiency led us to question whether rotenone-treated cells exhibit an iron starvation response despite iron accumulation.

To answer this question, we first probed the expression of proteins that respond to cytosolic labile iron status. We first quantified the expression of the transcription factor Hypoxia-inducible Factor 2-alpha (HIF2α), which is degraded through the ubiquitin-proteasome pathway after iron-mediated oxidation of the HIF2 protein via prolyl hydroxylase enzymes (26). Iron deficiency normally reduces HIF oxidation, thus preventing its degradation. Supporting cytosolic iron deficiency, rotenone increased HIF2α expression by immunoblot (Fig. 1F, 1H). The iron export protein ferroportin (FPN1) is normally downregulated in iron deficiency via translational repression with the iron-responsive proteins (IRPs) binding to an iron-responsive element (IRE) in the 5’- untranslated region (11, 21, 27). Consistent with this, we observed rotenone decreased FPN1 expression relative to vehicle controls (Fig. 1F, 1H). Cytosolic iron can be directly sensed by the O_2_ and iron sensor FBXL5, a master mammalian sensor of intracellular iron. Under iron replete conditions in neurons, FBXL5 normally binds iron by both an iron-binding hemerythrin-like N-terminal domain and a redox-sensitive [2Fe-2S] cluster near the C-terminus (23). Adequate oxygen levels maintain this ISC in a stable, oxidized state to activate FBXL5. Active FBXL5 recognizes and binds IRP2, which leads to IRP2 ubiquitination and degradation (23–25). Thus, FBXL5 in neurons under iron-replete conditions maintains IRP2 at low levels. Cytosolic iron deficiency prevents the proper FBXL5 sensing of iron, which causes FBXL5 instability and degradation. We predicted that cytosolic iron deficiency upon rotenone treatment would reduce FBXL5 levels. This was supported by western blot, in which we observed FBXL5 downregulation with rotenone treatment relative to controls (Fig. 1F, 1H).

Having observed changes in iron-responsive proteins consistent with cytosolic iron deficiency, we next asked whether reduced FBXL5 expression affected the IRP2 pathway. IRP2 is generally considered the dominant regulator of iron homeostasis in neurons than iron response protein 1 (IRP1) (27). IRP1 is primarily metabolic in neurons with a majority of IRP1 being in its cytosolic aconitase form that does not bind to IREs (38). Thus, IRP1 is secondary to IRP2 in neuronal iron regulation. Because of this, we investigated the protein expression of IRP2 and its targets. Treatment of SH-SY5Y neurons with excess iron-sulfate led to the expected changes associated with iron overload including IRP2 degradation and IRE-dependent reduction in *DMT1* and *TFR1*, but not *FTL1*, gene expression (Fig. S4). We next asked how rotenone affects IRP2 signaling. Consistent with functional iron deficiency in the cytosol, we observed that rotenone increased expression of IRP2 that was prevented by forcing iron overload with exogenous iron-sulfate treatment (Fig. 1G, 1H and Fig. S4B). However, in contrast to normal IRE regulation, we observed decreased expression of the iron import proteins TFR1 and DMT1 without changes in the FTH1 subunit of the iron storage protein ferritin (Fig. 1G, H). These results were further supported by observing a decrease in the *TFR1* and *DMT1* transcript levels by quantitative PCR (Fig. 1I). While the origin of this discrepancy remains unclear, the downregulation of iron import may partially induce the cytosolic labile iron deficiency we observed. Collectively, this data suggests perturbation of iron distribution and handling in mitochondria and cytosol.

### Rotenone induces ROS overproduction that mediates cell toxicity and death

Rotenone is a potent Complex I inhibitor that binds to the ubiquinone-binding (Q) site of Complex I (39, 40). This prevents the proper transfer of electrons originating from NADH oxidation. As a result, electron leakage from Complex I increases upon rotenone treatment to generate superoxide radicals. It is thus well accepted that rotenone induces cytotoxicity primarily through excess ROS production (41–44). We first confirmed rotenone treatment increased ROS levels using the fluorometric DCFDA assay. Consistent with increased ROS production, we observed rotenone treatment increased DCFDA fluorescence similar to H_2_O_2_ treated controls (Fig. 2A, 2B, and Fig. S5, S6). Increased ROS is partially neutralized via natural intracellular antioxidant defense and redox balance systems, including the oxidation of reduced glutathione (GSH) to the oxidized state (GSSG). Consistent with this, we observed that rotenone treatment promoted a shift in glutathione redox state towards the oxidized state with a lower GSH:GSSG ratio (Fig. 2C). Further, total glutathione species levels normally increase during extended oxidative stress through *de novo* glutathione synthesis from cysteine. We similarly observed adaptive increases in the cellular glutathione pool (Fig. 2D).

**Figure 2.**
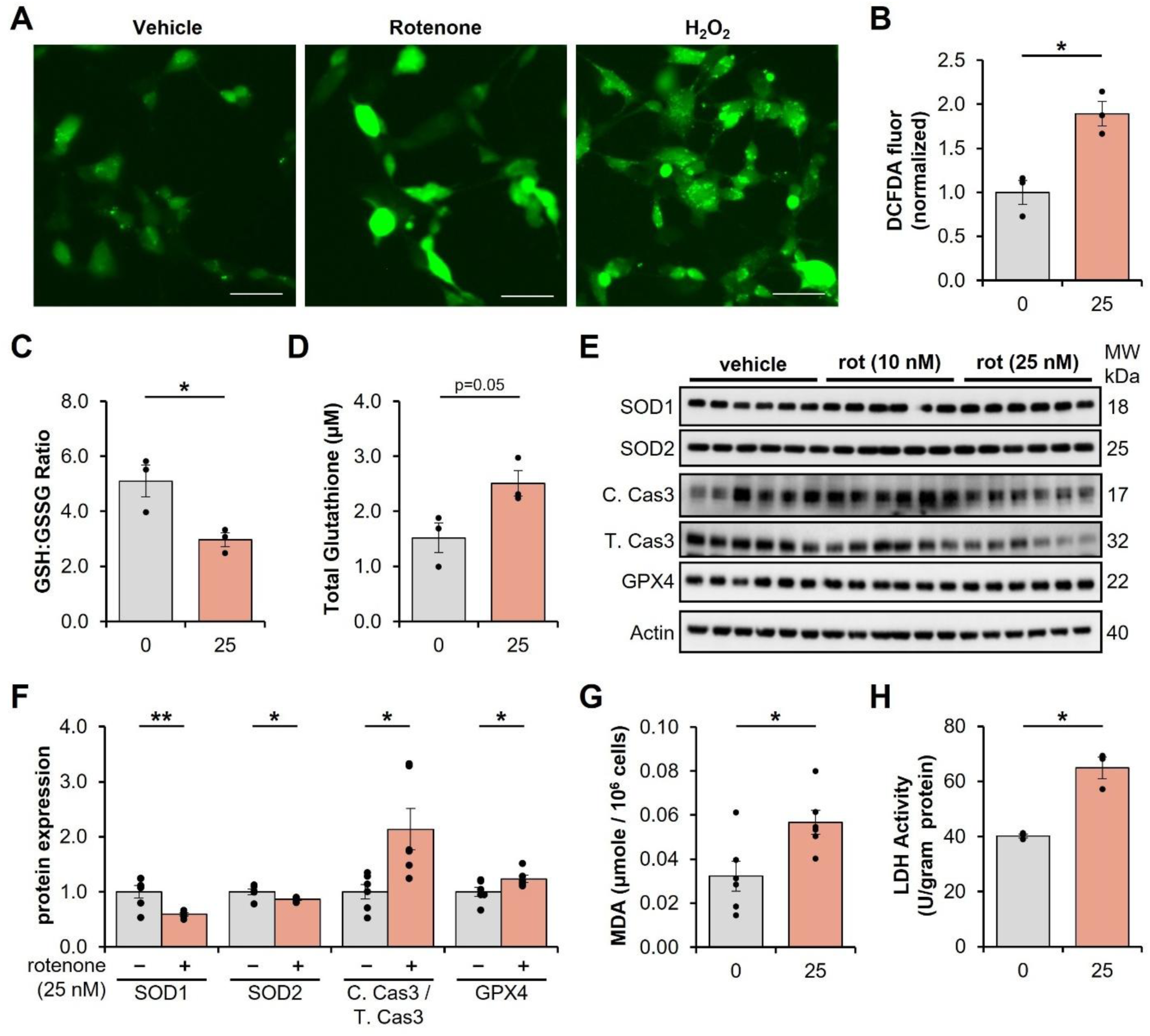
Rotenone causes oxidative damage and cell death in neurons. **(A)** Representative fluorescence microscopy images and **(B)** quantification (normalized to vehicle control) of DCFDA as a readout for ROS generation in differentiated SH-SY5Y cells treated with DMSO vehicle (0 nM) or rotenone (25 nM) for 24 hrs. Hydrogen peroxide (50 µM) was used as a positive control. Scale bars = 50 µm **(C)** Ratio of reduced (GSH) to oxidized (GSSG) glutathione levels and **(D)** quantification of total glutathione pool in differentiated SH-SY5Y cells treated with DMSO vehicle (0 nM) or rotenone (25 nM) for 24 hrs. **(E)** Representative images of western blot of proteins involved in cellular antioxidant defense (SOD1/2) and programmed cell death via cleavage (C. Cas3) of Procaspase 3 (T. Cas3) or ferroptosis (GPX4) in differentiated SH-SY5Y neurons treated with DMSO vehicle (0 nM) or rotenone (10 or 25 nM) for 24 hrs. **(F)** Densitometry of western blot images relative to actin or total protein and normalized to control levels for each individual protein in differentiated SH-SY5Y neurons treated with DMSO (0 nM) or rotenone (25 nM) for 24 hrs. C. Cas3/T. Cas3 represents the ratio of active Caspase-3 (C. Cas3) that was cleaved from the inactive Procaspase-3 (T. Cas3). **(G)** Quantification of MDA levels to assess lipid peroxidation via a colorimetric MDA-TBARS assay in differentiated SH-SY5Y neurons treated with DMSO vehicle (0 nM) or rotenone (25 nM) for 24 hrs. **(H)** Quantification of lactate dehydrogenase (LDH) activity in the media after exposure of differentiated SH-SY5Y neurons to DMSO vehicle (0 nM) or rotenone (25 nM) for 24 hrs. *p < 0.05, **p < 0.01

Under low and transient stress, glutathione homeostasis is sufficient to maintain the native cellular redox balance, however, severe ROS production and stress can overwhelm glutathione and antioxidant defense systems to damage biomolecules. Superoxide dismutase 1 (SOD1) in cytosol and SOD2 in mitochondria help neutralize harmful superoxide into less reactive hydrogen peroxide (45). Severe ROS production begins to damage these SOD1/2 proteins, leading to their degradation (45). Based on this mechanistic framework, we asked whether rotenone treatment decreased the abundance of SOD1 and SOD2. Consistent with severe ROS stress, we observed reduced SOD1 and SOD2 expression by immunoblot (Fig. 2E and 2F). We next asked whether rotenone is damaging other biomolecules, such as the oxidation of polyunsaturated fatty acids (PUFAs) and other lipids in the lipid membrane. We performed a colorimetric TBARS assay to quantify intracellular malondialdehyde (MDA), which is a breakdown byproduct of lipid peroxidation. Rotenone treatment nearly doubled intracellular MDA levels relative to control SH-SY5Y neurons (Fig. 2G). Thus, rotenone induces oxidative damage through ROS production.

We next asked whether rotenone reduces cell viability. Cytotoxic damage to the lipid membrane can cause leakage of lactate dehydrogenase (LDH) from cells into the surrounding media. We thus quantified LDH activity in vehicle- and rotenone-treated SH-SY5Y cells differentiated into dopaminergic neurons. LDH activity increased 50% in rotenone-treated cells, consistent with cytotoxicity (Fig. 2H). Rotenone has been reported to induce apoptosis, while iron-mediated lipid peroxidation is increasingly associated with ferroptosis in iron overload cells (29, 30, 46). We thus probed both apoptotic markers (e.g. cleaved Caspase-3) and ferroptotic markers (e.g. GPX4) to investigate whether rotenone is activating cell death pathways. Procaspase is cleaved to form caspase-3 (CAS3) upon induction of apoptosis, and GPX4 loss is a key trigger for ferroptosis. We thus quantified these markers by western blot. We observed an increase in cleaved CAS3 (Fig. 2E, 2F) and did not observe loss of GPX4 (Fig. 2E, 2F). Collectively, these data strongly suggest that rotenone treatment promotes oxidative stress and damage to induce apoptotic cell death.

### ROS drives iron misregulation through iron-sulfur cluster perturbations

Our overarching hypothesis suggests that the excess ROS generated by rotenone treatment in neurons causes the iron imbalances we observed. We thus sought to test this hypothesis directly by evaluating the changes in iron homeostasis in rotenone-treated cells upon quenching of ROS with antioxidants. To achieve this, we chose the well-established and potent brain-permeable antioxidant N-acetylcysteine amide (NACA), which replenishes glutathione to neutralize endogenous ROS (47). We first confirmed that NACA treatment (250 µM) concomitantly with rotenone in SH-SY5Y differentiated neurons prevents excess ROS generation relative to vehicle-treated cells using a DCFDA fluorescent assay (Fig. 3A, 3B and Fig. S6). As expected, we observed increased fluorescence, consistent with increased ROS, in the rotenone-treated cells (Fig. 3A and 3B). Treatment with the antioxidant NACA prevented excess ROS generation (Fig. 3A and 3B). Excess iron accumulation enhances ROS production through the Fenton reaction. We asked whether the excess iron upon rotenone treatment contributes to the ROS production and perpetuates a self-propagating cycle of ROS generation. To do this, we treated rotenone-treated cells with the iron chelator deferoxamine (DFO, 25 µM) and evaluated ROS levels by DCFDA assay. We observed DFO prevented excess ROS generation (Fig. 3A, 3B and Fig. S6), consistent with excess iron inducing oxidative stress. However, these data do not definitively establish whether ROS is the cause of iron misregulation.

**Figure 3.**
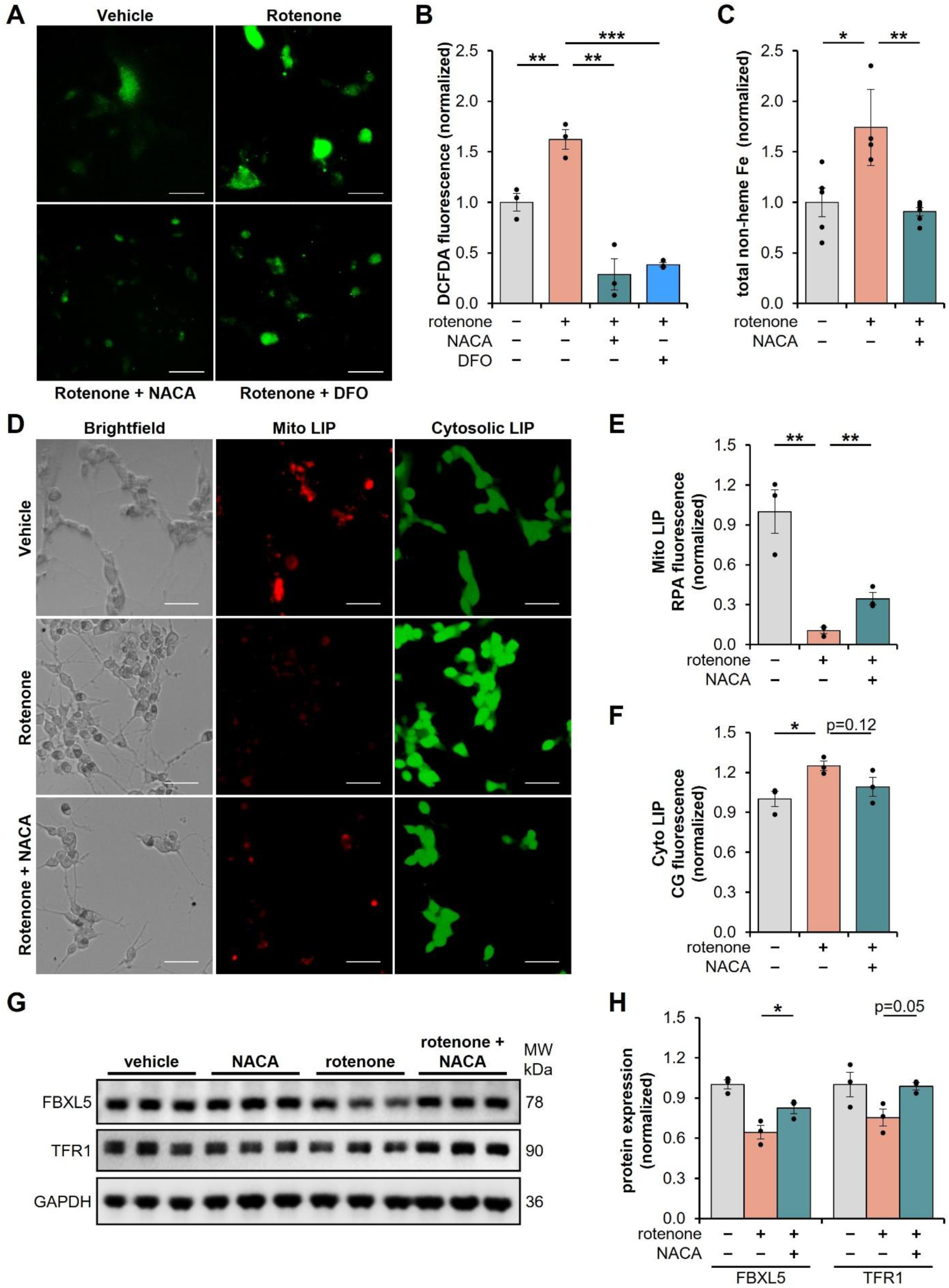
ROS generation drives iron misregulation upon rotenone treatment in neurons. **(A)** Representative fluorescence microscopy images and **(B)** quantification (normalized to vehicle control) of DCFDA as a readout for ROS generation in differentiated SH-SY5Y cells treated with DMSO vehicle (0 nM) or rotenone (25 nM) for 24 hrs. Some cells were also simultaneously treated with either the antioxidant N-acetylcysteine amide (NACA, 250 µM) or deferoxamine (DFO, 25 µM) in the presence of rotenone (25 nM) for 24 hrs. **(C)** Quantification of total cellular non-heme iron levels via a colorimetric ferrozine assay in differentiated SH-SY5Y neurons treated with DMSO vehicle (0 nM), rotenone (25 nM), or NACA (250 µM) for 24 hrs. **(D)** Representative images and **(E**, **F)** quantification of labile iron pools in the mitochondria and cytosol in differentiated SH-SY5Y cells treated with rotenone (25 nM) or DMSO vehicle (0 nM) using the iron-responsive fluorescent dyes RPA (mitochondria, red) and Calcien Green-AM (cytosol, green). Scale bars = 50 µm. **(G)** Representative western blot images and **(H)** densitometric analysis of proteins involved in iron homeostasis in differentiated SH-SY5Y neurons treated with DMSO vehicle, rotenone (25 nM), or NACA (250 µM) in the presence of rotenone (25 nM) for 24 hrs. Protein was quantified relative to GAPDH and normalized to control for each individual protein. *p < 0.05, **p < 0.01, ***p < 0.001

To test our hypothesis that ROS is driving iron misregulation, we next asked whether antioxidant treatment, which prevents excess ROS generation, is sufficient to restore normal iron balance in rotenone-treated neurons. Consistent with our hypothesis, non-heme iron levels increased in rotenone-treated cells (Fig. 3C), however, antioxidant treatment with NACA in the presence of rotenone prevented iron accumulation (Fig. 3C). We next asked whether NACA was able to prevent imbalances in intracellular iron distribution using iron-responsive fluorescent dyes RPA (for mitochondrial LIP) and calcein green-AM (for cytosolic LIP). Similar to our previous experiments, we observed that rotenone treatment quenched RPA fluorescence in mitochondria, consistent with increased mitochondrial labile iron (Fig. 3D, 3E and S3). We also observed increases in calcein green fluorescence upon rotenone treatment relative to vehicle controls, consistent with cytosolic iron deficiency (Fig. 3D, 3F and S3). Antioxidant treatment with NACA partially reversed these effects (Fig. 3D, 3E and S3). These results suggest that ROS is causative in inducing iron maldistribution, and suggests antioxidant treatment may restore normal expression and function of iron-dependent proteins. We thus quantified expression of TFR1 and FBXL5 by immunoblot. Consistent with the capacity for antioxidant treatment to prevent iron maldistribution in rotenone-treated cells, we similarly observed restored expressions of both FBXL5 and TFR1 in neurons treated with both rotenone and NACA (Fig. 3F and 3G). Collectively, this data suggests ROS induces iron overload, which in turn stimulates further ROS production, creating a cycle that persists until either ROS or iron is neutralized in the system.

Our results thus led us to question how ROS production causes iron misregulation. We observed that mitochondria are especially sensitive to iron imbalances upon rotenone treatment. As rotenone inhibits Complex I leading to ROS leakage in mitochondria, we first explored pathways that utilize mitochondrial iron. Most mitochondrial iron is either incorporated into proteins such as the electron transport chain, utilized in heme synthesis, or is involved in ISC biogenesis and utilization (13–15, 48). ISCs are primarily found in either the often more sensitive [4Fe-4S] or the resistant [2Fe-2S] states (49, 50). Loss of ISC protein integrity, such as damage from ROS-mediated oxidation, frequently cause structural changes that induce clearance via mitochondrial proteases or ubiquitination and subsequent degradation of the corresponding ISC-containing protein (51–53). We thus quantified expression of proteins that contain either [4Fe-4S] or [2Fe-2S] clusters in the absence or presence of rotenone.

NDUFAB is a highly soluble accessory subunit adjacent to the rotenone-binding Q-module on Complex I (Fig. 4A) and is critical for ISC regulation (54). However, NDUFAB does not contain an ISC *per se* and was thus used as a control. We predicted as it does not contain an ISC and is stable to mitochondrial Lon protease, it would be resistant to rotenone-mediated ROS generation. As expected, we did not observe any changes in NDUFAB expression upon rotenone treatment by immunoblot (Fig. 4B, C). Consistent with our hypothesis, rotenone reduced expression of NDUFS1 (Fig. 4A-C), which is the largest of 45 Complex I subunits and contains the three ISCs N1b ([2Fe-2S]), N4 ([4Fe-4S]), and N5 ([4Fe-4S]) (39, 40). Further, rotenone reduced expression of the accessory subunit NDUFS4 (Fig. 4B, C), which is located near multiple ISCs (Fig. 4A) and plays a critical role in the mature assembly of Complex I (55). We next questioned whether rotenone-mediated ROS generation affects other ISC-containing proteins in mitochondria and the cytosol. Ferrochelatase (FECH) is the terminal enzyme in heme biosynthesis, which forms a matrix-facing functional homodimer in the inner mitochondrial membrane and contains an oxidative-sensitive [2Fe-2S] ISC (56). We observed a non-significant trend in the reduction of FECH expression (Fig. 4B, C). Finally, mitochondrial and cytosolic aconitase both contain an [4Fe-4S] ISC that is required for their enzymatic activity. Rotenone treatment decreased the activity of mitochondrial aconitase (Fig. 4D), however, had no effect on cytosolic aconitase activity (Fig. 4D). Collectively, this data suggests rotenone-mediated ROS generation in mitochondria affects ISC protein dynamics in mitochondria, but not in cytosol.

**Figure 4.**
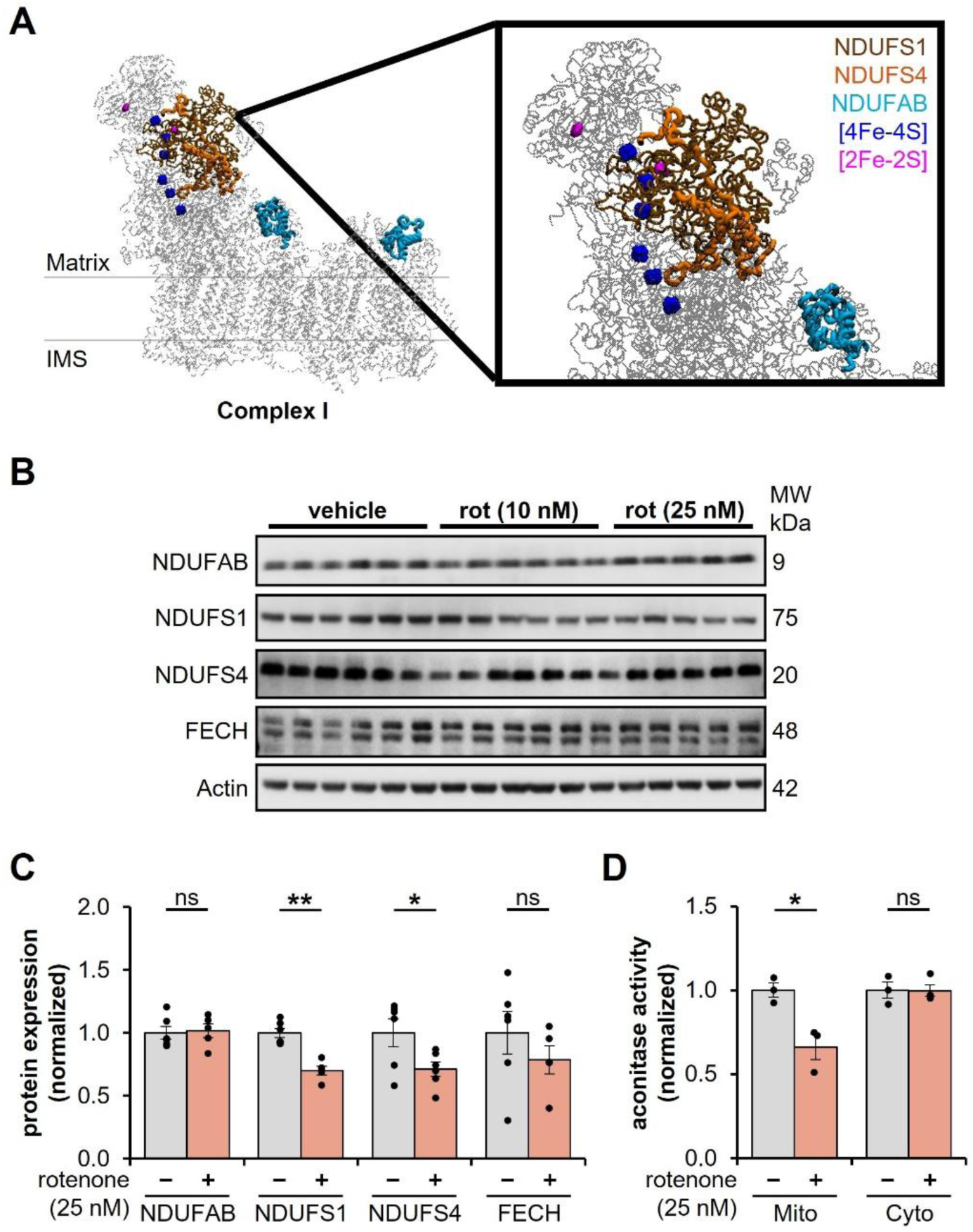
ROS alter iron-sulfur cluster dynamics to induce iron dyshomeostasis. **(A)** Cryo-EM structure (PDB 5XTD) from human cardiac cells of specific Complex I subunits and iron-sulfur clusters. **(B)** Representative western blot images and **(C)** densitometric analysis of Complex I subunits and the ISC-containing protein ferrochelatase (FECH) in differentiated SH-SY5Y neurons treated with DMSO vehicle or rotenone (25 nM) for 24 hrs. Protein was quantified relative to Actin and normalized to control for each individual protein. **(D)** Aconitase activity in isolated mitochondrial and cytosolic fractions (normalized to control) in differentiate SH-SY5Y neurons treated with DMSO vehicle or rotenone (25 nM) for 24 hrs. *p < 0.05, **p < 0.01, ns = not significant

## Discussion

Mitochondrial Complex I dysfunction and iron accumulation are well-established hallmarks of Parkinson’s disease, yet the mechanistic sequence connecting these abnormalities in dopaminergic neuronal cell death in the substantia nigra remains unclear. It is increasingly recognized that Complex I deactivation, through either environmental or genetic mechanisms, induces progressive parkinsonism features (2, 6). Here, we demonstrate that inhibition of Complex I by rotenone initiates a ROS-dependent redistribution of intracellular iron in SH-SY5Y cells differentiated into dopaminergic neurons. Rotenone increased mitochondrial and total cellular iron while simultaneously decreasing the cytosolic labile iron pool, revealing that cellular iron accumulation can coincide with iron starvation in the cytosol. Antioxidant treatment prevented these changes, indicating that ROS generation is required for the disruption of iron homeostasis. Conversely, iron chelation reduced ROS, supporting a positive feedback mechanism in which Complex I-derived ROS initially perturb iron handling, and the resulting mitochondrial iron accumulation subsequently amplifies oxidative stress through Fenton chemistry or other mechanisms. Our findings therefore provide a mechanistic link between two central hallmarks of PD by placing mitochondrial ROS upstream of iron maldistribution (Fig. 5). This model expands upon earlier work by Sherer and colleagues demonstrating that rotenone toxicity is driven primarily by oxidative damage downstream of Complex I inhibition rather than ATP depletion alone (41, 44). Together, these findings suggest that iron accumulation in PD may not simply represent a late consequence of neuronal injury but may arise from an early failure to coordinate mitochondrial and cytosolic iron utilization during oxidative stress.

**Figure 5.**
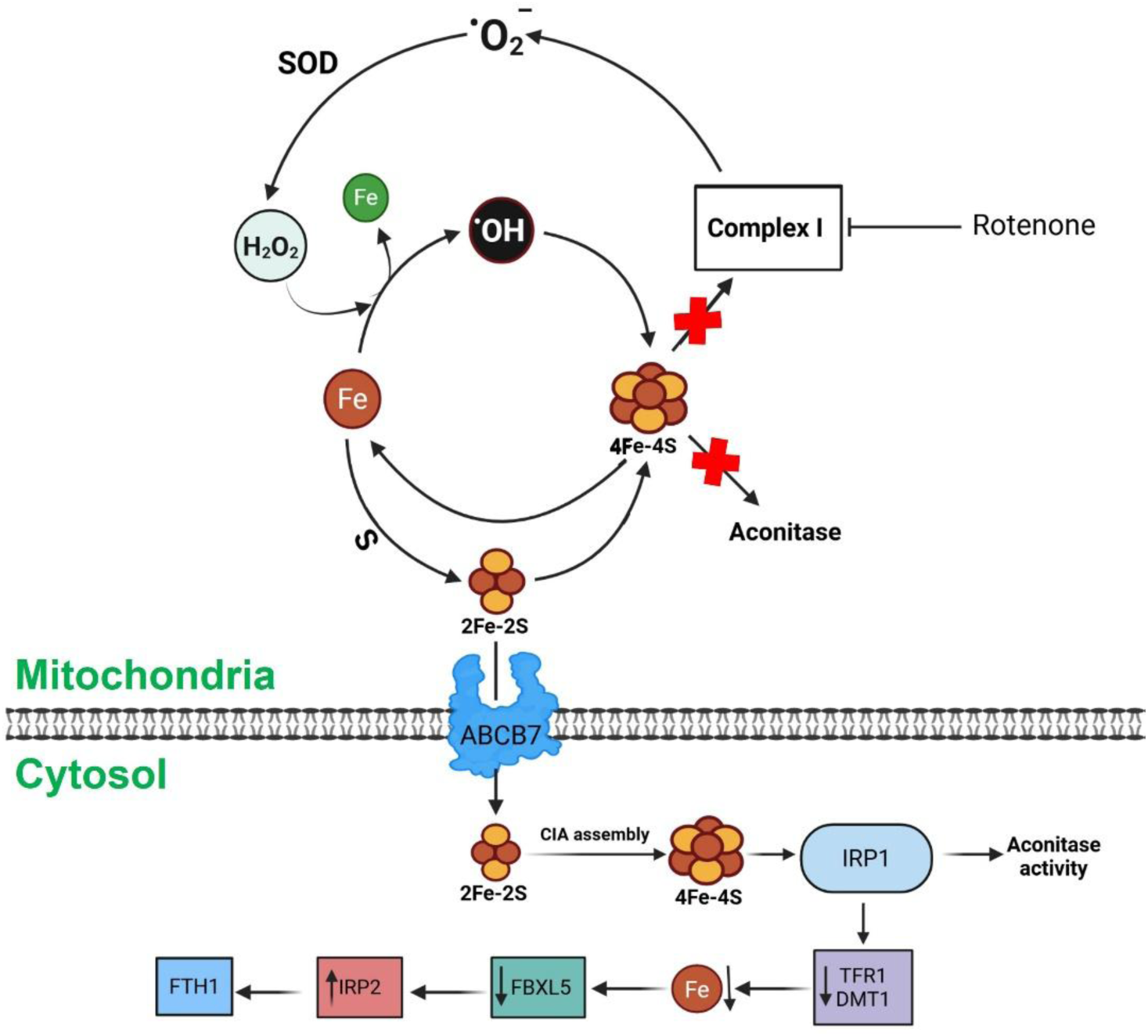
Simplified schematic of iron misregulation through excess ROS production with rotenone treatment. Created in BioRender. Lab, G. (2026) https://BioRender.com/hm4yhol. We propose inhibition of Complex I with rotenone induces an increase in superoxide that is converted to hydrogen peroxide via SOD1/2. In the presence of mitochondrial labile iron, the excess hydrogen peroxide undergoes the Fenton reaction, producing hydroxyl radicals, which affects [4Fe-4S] clusters, leading to an increase in the mitochondrial LIP, and [2Fe-2S] clusters to induce the misregulation of the cytosolic iron-related proteins. The perturbations in mitochondrial [4Fe-4S] clusters affect the structure and function of ISC-dependent proteins, such as Complex I and mitochondrial aconitase enzyme activity.

Rotenone is a particularly informative and leading model for examining this relationship because it reproduces several biochemical and pathological features associated with PD, including selective dopaminergic neuron injury, oxidative damage, mitochondrial dysfunction, and α-synuclein accumulation (9). Epidemiological and experimental studies have also associated pesticide exposure, including rotenone exposure, with increased PD risk, although exposure alone should not be interpreted as sufficient to cause every case of PD. The mechanisms identified here may extend to other neurotoxins that converge upon mitochondrial dysfunction and ROS production. 1-methyl-4-phenyl-1,2,3,6-tetrahydropyridine (MPTP) is converted to MPP+, which accumulates in dopaminergic neurons and deactivates Complex I (9), while 6-hydroxydopamine (6-OHDA) undergoes oxidation to generate superoxide, hydrogen peroxide, hydroxyl radicals, and quinone species. Lee *et al.* further showed that 6-OHDA induces iron accumulation, ROS generation, and lipid peroxidation in SH-SY5Y cells (57), suggesting that disruption of iron homeostasis may be shared across multiple environmental toxin-based PD models. Thus, although the initiating event differs among rotenone, MPTP, and 6-OHDA, mitochondrial oxidative stress may represent a common event that destabilizes iron-containing proteins and promotes iron retention in vulnerable dopaminergic neurons.

A central finding of our study is that rotenone-treated neurons respond as if they are iron deficient despite accumulating iron at the cellular and mitochondrial levels. Reduced cytosolic labile iron was accompanied by increased HIF2α, decreased FBXL5, stabilization of IRP2, and reduced FPN1, which are collectively consistent with activation of a cytosolic iron-starvation response. The stabilization of IRP2 is particularly important because FBXL5 ordinarily promotes IRP2 degradation under iron-replete conditions. However, the responses of some downstream IRE-regulated proteins did not follow the canonical model. Rather than increasing TFR1 and DMT1 to promote iron uptake, rotenone reduced their protein and transcript levels, while FTH1 abundance was unchanged. These results suggest that the IRE/IRP system is activated but is not operating in isolation, or that this was the cause of cytosolic iron starvation. However, mitochondrial iron accumulation suggests alternative pathways. Oxidative stress-dependent transcriptional regulation, altered transcript stability, impaired protein synthesis, or compensatory mechanisms that limit further iron uptake may supersede the expected effects of IRP2 on individual targets.

Importantly, N-acetylcysteine amide (NACA) restored FBXL5 and TFR1 expression while correcting the mitochondrial and cytosolic labile iron pools. This indicates that the atypical IRE/IRP response is not simply an unrelated consequence of rotenone toxicity, but is coupled to the ROS-dependent redistribution of iron. Our results therefore emphasize that total cellular iron measurements alone cannot define the iron status experienced by individual organelles or iron-sensing pathways.

Our data further identify mitochondrial ISC proteins as potential molecular targets linking ROS production to iron maldistribution. ISCs are intrinsically redox active, and their oxidation can impair enzymatic activity, destabilize the surrounding protein, and release iron that may further contribute to the mitochondrial labile iron pool. The selective loss of NDUFS1 and mitochondrial aconitase activity, together with the relative preservation of ferrochelatase and cytosolic aconitase activity, supports a spatially restricted effect that is strongest near the mitochondrial source of rotenone-induced ROS generation. NDUFS1 is especially relevant because it coordinates one [2Fe-2S] cluster and two [4Fe-4S] clusters within the N-module of Complex I. By contrast, ferrochelatase contains a [2Fe-2S] cluster and exhibited only a nonsignificant trend toward reduced abundance. Although individual cluster sensitivity is influenced by solvent accessibility, ligation, protein environment, and local ROS exposure, the collective results are consistent with preferential disruption of sensitive mitochondrial [4Fe-4S]-containing proteins. This interpretation is also supported by the loss of mitochondrial aconitase activity, which requires an intact [4Fe-4S] cluster, without a corresponding loss of cytosolic aconitase activity. Mitochondrial [4Fe-4S] cluster maturation requires a later and more complex pathway involving reductive fusion of [2Fe-2S] precursors, whereas transfer to mitochondrial [2Fe-2S] target proteins can occur more directly (48). Oxidative damage may therefore simultaneously destroy existing [4Fe-4S] clusters and create an increased demand for a comparatively complex biosynthetic pathway, promoting ineffective iron reutilization and mitochondrial iron retention. Thus, the transport of an ISC-related metabolite from the mitochondria to the cytosol via ABCB7 may contribute to the perturbed cytosolic labile iron response (48).

The differential behavior of NDUFS1, NDUFS4, and NDUFAB provides additional insight into how cluster damage may affect Complex I structure. NDUFS1 contains three ISCs and is positioned within the N-module, whereas NDUFS4 is an accessory subunit required for the stability and maturation of the intact complex (Fig. 4A). NDUFAB does not contain an ISC and remains stable following rotenone treatment. Oxidation or loss of the NDUFS1 clusters may destabilize local protein folding, expose normally protected regions, and promote degradation of damaged NDUFS1. Loss of NDUFS1 could then alter the architecture of the N-module and weaken its association with the remainder of Complex I. NDUFS4 may be lost as part of this secondary structural destabilization even though it does not directly coordinate an ISC. Consistent with this possibility, cryo-EM structural analyses of mouse heart Ndufs4-deficient Complex I revealed a weak association of the N-module in Complex I missing NDUFS4, supporting an important role for NDUFS4 in maintaining N-module integration (55). NDUFS1 and NDUFS4 may also be preferentially recognized by the mitochondrial Lon protease after oxidative unfolding or Complex I disassembly (58), whereas NDUFAB may be comparatively resistant because it is not located on the N-module and as a result, is not similarly exposed as a Lon-sensitive substrate. Thus, the preservation of NDUFAB does not establish that the entire N-module remains intact. Instead, it suggests selective proteolysis of damaged or structurally exposed components rather than indiscriminate degradation of Complex I. This model predicts that rotenone should decrease intact N-module or holo-Complex I assembly and that inhibition or depletion of LONP1 should partially preserve NDUFS1 and NDUFS4.

Finally, our findings help distinguish the initiating mechanism of iron misregulation from the downstream mode of cell death. Rotenone increased ROS, oxidized the glutathione pool, decreased SOD1 and SOD2, increased lipid peroxidation, and caused loss of membrane integrity. These features overlap with ferroptosis, which is characterized by iron-dependent lipid peroxidation and is increasingly implicated in PD. Recent work has reported ferroptosis-associated changes, including lipid peroxidation and GPX4-ACSL4 dysregulation, in rotenone-treated SH-SY5Y cells (46). However, our experimental conditions increased cleaved caspase-3 without reducing GPX4, supporting apoptosis as the predominant detectable cell-death pathway. These outcomes are not necessarily contradictory. Iron-dependent ROS propagation and lipid damage can contribute to cellular injury without achieving initiation of ferroptosis. The dominant mode of death may also depend upon rotenone concentration, exposure time, differentiation state, antioxidant capacity, and the severity of mitochondrial damage. Accordingly, our data support a model in which ROS-mediated ISC damage and iron redistribution precede apoptotic death, while creating intracellular conditions that could increase susceptibility to ferroptosis under prolonged or more severe stress.

More broadly, age-related neurodegenerative diseases arise from interconnected dysfunction of mitochondrial activity, metal homeostasis, proteostasis, and antioxidant defense rather than disruption of a single pathway. Defining the cellular sequence that connects these processes is essential for identifying interventions capable of intervening in disease progression. A more fundamental understanding of how reduced Complex I activity alters ISC integrity and intracellular iron distribution may ultimately guide therapeutic strategies for PD, Leigh syndrome, and related disorders characterized by mitochondrial dysfunction and progressive neuronal loss.

## Experimental Procedures

### Statistical analysis

Data, from a minimum of three independent biological replicates, are presented as weighted means ± SEM, with multiple technical replicates included where applicable unless otherwise noted. Samples and data sets were randomized, and analyses were conducted independently by multiple participants whenever possible. P-values for the experiments were calculated using a two-tailed Student’s T-test or ANOVA with multiple comparisons where appropriate. * p≤0.05, ** p<0.01, *** p<0.001, **** p<0.0001, ns=Not Significant.

### Cell culture and cell differentiation

SH-SY5Y cells (ATCC CRL-2266) were cultured in DMEM Dulbecco’s Modified Eagle’s Medium, SH30022.FS) supplemented with 10% FBS, 1% glutamine, 1% penicillin-streptomycin, and 3.7 g/L sodium bicarbonate at 37 °C with 5% CO_2_ and controlled humidity. Cells were differentiated into dopaminergic neurons according to established procedures (59), with a slight modification of using 10% FBS. Unless otherwise specified, culture medium was aspirated from differentiated SH-SY5Y cells and replaced with fresh medium containing vehicle (DMSO, 0.1%), rotenone (10 or 25 nM in DMSO from 1000X stock), N-acetylcysteine amide (250 µM in DMSO from 1000X stock), or iron sulfate (250 µM in water from 100X stock), and cells were incubated for 24 hours.

### Mitochondrial isolation

Mitochondria were isolated from the corresponding differentiated SH-SY5Y cell line, after harvesting and counting the cells using the Mitochondria Isolation Kit for Cultured Cells (Abcam, Cat. # ab110170), following the manufacturer’s protocol.

### Quantification of non-heme iron

A modified ferrozine-based assay was used to quantify levels of non-heme iron in differentiated SH-SY5Y cells and isolated mitochondria. Samples were digested with 3M HCl containing 600 mM trichloroacetic acid (TCA). Samples were heated at 95 °C for 4 hours, centrifuged, and the supernatant collected. Chromogen solution (100 µL containing 100 mM ferrozine, 1.5 M sodium acetate, 1.5% v/v thioglycolic acid in deionized water) was added to the digestate (50 µL) in a 96-well plate and incubated for 45 minutes at 37 °C to form a magenta-colored solution. Absorbance was measured with three technical replicates by a microplate reader at 562 nm in a 96-well plate and quantified using a standard curve. Data was normalized based on cell number and mitochondrial protein concentration, respectively. A standard curve was prepared from a freshly prepared 1 g/L iron stock (10 mL containing 18 mM ferrous sulfate heptahydrate and 32 mM ascorbic acid in deionized water) followed by serial dilutions to a final concentration of 0, 0.1, 0.5, 1, 2.5, 5, 10, and 20 mg/L.

### Assessment of lipid peroxidation

Lipid peroxidation was quantified using a TBARS Assay Kit (Cayman Chemicals 10009055) similar to the manufacturer’s protocol with slight modifications. Cell pellets were homogenized and lysed using PBS. Cell debris was removed by centrifugation, and protein concentration was calculated from the resulting supernatant. MDA-TBA adduct was prepared according to the manufacturer’s protocol, except at 2X concentrations to increase signal. MDA-TBA adduct was quantified using a standard curve after reading absorbance at 530 nm with a plate reader in a 96-well plate.

### Protein analysis via Western Blot

Proteins were extracted from the cells with RIPA buffer containing protease and phosphatase inhibitors. After centrifugation, protein extract was collected, and protein concentration was quantified by a BCA assay. Samples for Western blot were prepared by diluting protein extract (∼20 µg) in Laemmli sample buffer and reducing agent. Samples were denatured at 95 °C for 5 minutes and loaded onto Bolt™ 4-12% MIDI or MINI gels. Proteins were separated by electrophoresis at 150 V in MOPS running buffer. Protein was transferred to a PVDF membrane using a Trans-Blot® Turbo Transfer system with the manufacturer’s transfer buffer containing an additional 0.2% SDS. Total protein was imaged according to the manufacturer’s protocol (Thermo Fisher A44717), and blots were blocked with 5% BSA in TBST at room temperature for 30 minutes. Membranes were incubated in 5% BSA with primary antibodies overnight at 4 °C, followed by 1 hour at room temperature. Membranes were rinsed with TBST (3 x 10 minutes) and incubated with secondary antibody for 1-2 hours in 5% BSA in TBST. After rinsing membranes with TBST, chemiluminescence was detected using a chemiluminescence kit (Thermo Fisher SuperSignal PicoPlus) and imaged with an iBright FL1500 imager. Membranes were stripped for 5 minutes with stripping buffer, rinsed with TBST, re-blocked, and re-probed as described above. Densitometry was performed using iBright Analysis Software or ImageJ analysis and quantified relative to actin loading control or total protein (imaged using total protein stain). Relative protein was normalized to control levels. A list of antibodies and dilutions can be found in Table 1.

**Table 1.** Western Blot Antibody Information.

| **Name** | **Company** | **Identifier** | **Dilution** |
| --- | --- | --- | --- |
| Anti-TFR1 | Cell Signaling Technology | 13113S | 1:10,000 |
| Anti-DMT1 | Santa Cruz Biotechnology | sc-166884 | 1: 3,000 |
| Anti-FTH1 | Cell Signaling Technology | 3998S | 1:1,000 |
| Anti-IRP2 | Cell Signaling Technology | 37135S | 1:1,000 |
| Anti-FBXL5 | Thermo Fisher Scientific | PA5-113529 | 1:3,000 |
| Anti-SOD1 | Cell Signaling Technology | 2770S | 1:3,000 |
| Anti-SOD2 | Santa Cruz Biotechnology | sc-133134 | 1:3,000 |
| Anti-NDUFS1 | Cell Signaling Technology | 70264S | 1:1,000 |
| Anti-NDUFS4 | Santa Cruz Biotechnology | sc-100567 | 1:1,000 |
| Anti-Cleaved<br>Caspase 3 | Cell Signaling Technology | 9661T | 1,1000 |
| Anti-Total<br>Caspase 3 | Cell Signaling Technology | 9662S | 1,1000 |
| Anti-GPX4 | Cell Signaling Technology | 52455S | 1:1,000 |
| Anti-FECH | GeneTex | GTX65973 | 1:3,000 |
| Anti-HIF2α | Cell Signaling Technology | 7096S | 1:3,000 |
| Anti-TH | Cell Signaling Technology | 2792S | 1:500 |
| Anti-MAP2 | Cell Signaling Technology | 4542S | 1:500 |
| Anti-Actin HRP<br>Conjugate | Cell Signaling Technology | 5125S | 1:20,000 |
| Anti-GAPDH<br>HRP Conjugate | Cell Signaling Technology | 3683S | 1:10,000 |
| Donkey Anti-Rabbit HRP<br>Conjugate | Thermo Fisher Scientific | 31458 | 1:20,000 |
| Horse Anti-Mouse HRP<br>Conjugate | Cell Signaling Technology | 7076S | 1:5,000 |
| Donkey Anti-Rabbit, Alexa<br>Fluor 488 | Thermo Fisher Scientific | A-21206 | 1:1,000 |
| No Stain Protein<br>Labeling<br>Reagent | Thermo Fisher Scientific | A44449 | - |

### mRNA quantification via qPCR

Total RNA was isolated from cells according to manufacturer protocols (Thermo Fisher PureLink RNA Mini Kit) and quantified with a NanoDrop spectrophotometer. Primers were designed and checked by NCBI Primer BLAST for sequence specificity and potential cross-reactivity. Primers used can be found in Table 2. Primers (IDT) were diluted to 1 mM stocks. Relative mRNA expression was quantified by qRT-PCR using a One-Step RT-PCR Kit (Biorad iTaq Universal SYBR Green) containing 100 nM primer and 150 ng total RNA in a 10 μL reaction. PCR reaction products were verified by melting temperature. Gene expression was quantified using the Pfaffl Method relative to *GAPDH* expression and normalized to control group levels.

**Table 2.**
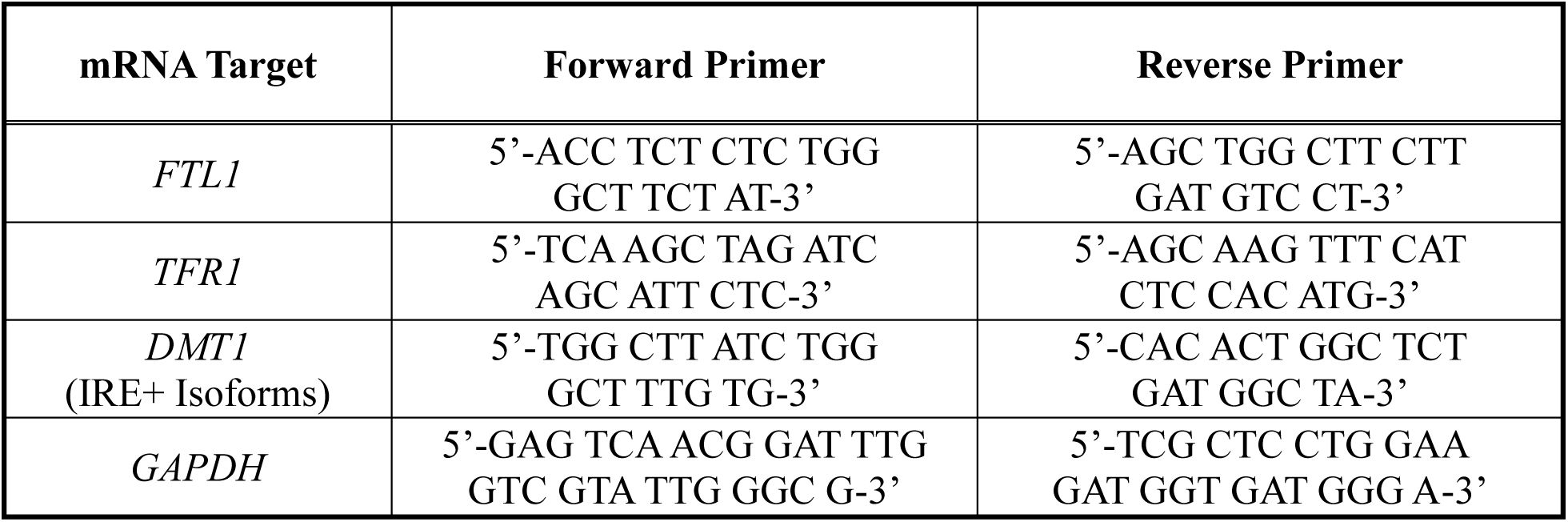
Primers used for qPCR.

### Visualization of labile iron

Differentiated cells were treated with vehicle, rotenone, and/or antioxidant for 24 hours. Media was then changed with fresh media containing calcein green-AM (8 μM, AAT Bioquest 22002) or RPA (10 μM, Axxora SQX-RPA.1) from 1000X stocks in DMSO. Cells were incubated for 30 minutes, media removed, cells rinsed with PBS, and fluorescence imaged via a digital fluorescence microscope or confocal microscope. Relative fluorescence per cell was quantified via ImageJ in a minimum of three independent images per well (>100 cells per biological replicate).

### Glutathione ratio

Reduced to oxidized glutathione was quantified in differentiated SH-SY5Y neurons using a glutathione assay kit (Abcam ab205811) according to manufacturer protocols. Cells were collected at ∼4x10^5^ cells/well and washed with PBS. Cells were removed mechanically and resuspended in 100 μL of cold PBS containing 0.5% NP-40 buffer. Cell lysate was centrifuged at 1000 rcf at 4 °C for 15 minutes. Supernatant was collected and deproteinized using a pH=5-6 buffer. After incubating for 30-45 minutes in the dark at room temperature, fluorescence intensity (Ex/Em = 490 nm/520 nm) was measured using a Microplate Reader with technical duplicates.

### ROS visualization

Relative ROS generation was quantified using a DCFDA Cellular ROS Detection Assay Kit (Abcam ab113851) according to manufacturer protocols. After 24 hours of treatment, media was aspirated and cells rinsed with warm PBS. After aspiration, DCFDA (1000X stock in DMSO) was added to dilution buffer (20 μM final concentration) and added to the cells. Fluorescence was imaged on a digital fluorescent microscope with a minimum of three images per well. Relative DCFDA fluorescence was then quantified using ImageJ and normalized to cell count. Hydrogen peroxide (50 μM) was used as a positive control.

### Lactate dehydrogenase activity

Differentiated cells were treated with either vehicle or rotenone for 24 hours. Supernatant of cultured rotenone-treated and control neurons was collected to determine lactic dehydrogenase (LDH) activity using the commercial LDH activity assay kit (Elabscience Biotechnology). The activity of LDH was determined following the manufacturer’s guidelines.

### Aconitase activity

Mitochondrial and cytosolic aconitase activity was determined using the Aconitase Activity microplate assay kit (Abcam ab109712) by colorimetrically monitoring the conversion of isocitrate to *cis*-aconitate at 240 nm according to manufacturer protocols.

### Immunofluorescence of cell differentiation

Sterilized glass coverslips were positioned in the wells of a 24-well plate prior to cell seeding. After following the differentiation protocol for the cells, the medium was discarded, washed 3 times with PBS, and the cells were fixed with 4% paraformaldehyde diluted with PBS at 37 °C for 20 minutes. After washing with PBS three times, the cells were permeabilized in 0.2% Triton X-100 at room temperature for 10 minutes, blocked with 3% BSA in PBS for 1 hour, and incubated with either Tyrosine Hydroxylase or MAP2 primary antibody overnight at 4 °C. Then, the corresponding fluorescent secondary antibody was used at room temperature for 1 hour. The samples were first washed three times with PBS. DAPI, diluted to 2 μg/mL in the blocking solution, was then added, and the samples were incubated in the dark for 5 minutes. The images were captured with a ZOE fluorescence microscope. The colocalization between two fluorescence signals in cells was quantified by ImageJ software.

## Data Availability

Raw data used in this work is available in supplemental data. Any relevant data in addition to that included in this manuscript will be provided upon reasonable request or made available publicly (e.g. Dryad).

## Supporting Information

This article contains supporting information. Uncropped and unedited immunoblot images can be found in the Supplemental Data.

## Acknowledgements

We thank the MSU Veterinary Diagnostics Laboratory for performing ICP-MS determination of total iron levels. We acknowledge the use of Visual Molecular Dynamics (VMD) for structure visualization of Complex l with assistance from Maryum Irshad. We thank Mayowa Ojumah & Sushila Thapa with assistance on cell maintenance.

## Author Contributions

ASAH and ASG conceived the project and developed the overall concept. ASG provided funding. ASAH, SP, KHP, MAM, and ARA performed the experiments. ASAH and ASG analyzed the data. ASAH and ASG contributed important intellectual input. ASAH and ASG wrote and edited the manuscript.

## Funding and additional information

This work was generously supported by the University of Cincinnati, the United Mitochondrial Disease Foundation (PI-23-0026), and the National Institute of Health National Institute on Aging (1R01AG084718). ARA was supported by an NSF-REU Fellowship (CHE-1950244). KHP was supported by the UC McNair Scholars Program.

## Abbreviations and nomenclature

ABCB7: ATP-binding cassette subfamily B member 7
ACSL4: acyl-CoA synthetase long-chain family member 4
ANOVA: analysis of variance
BCA: bicinchoninic acid
BSA: bovine serum albumin
CAS3: caspase-3
CI: complex I
DAPI: 4′,6-diamidino-2-phenylindole
DCFDA: 2′,7′-dichlorodihydrofluorescein diacetate
DFO: deferoxamine
DMEM: Dulbecco’s modified Eagle’s medium
DMSO: dimethyl sulfoxide
DMT1: divalent metal transporter 1
ETC: electron transport chain
FBXL5: F-box and leucine-rich repeat protein 5
FECH: ferrochelatase
FBS: fetal bovine serum
FPN1: ferroportin 1
FTH1: ferritin heavy chain 1
FTL1: ferritin light chain 1
GPX4: glutathione peroxidase 4
GSH: reduced glutathione
GSSG: oxidized glutathione
HIF1α: hypoxia-inducible factor 1 alpha
HIF2α: hypoxia-inducible factor 2 alpha
HRS: hours
ICP-MS: inductively coupled plasma-mass spectrometry
IRE: iron-responsive element
IRP1: iron regulatory protein 1
IRP2: iron regulatory protein 2
ISC: iron-sulfur cluster
LDH: lactate dehydrogenase
LIP: labile iron pool
LONP1: Lon peptidase 1, mitochondrial
MAP2: microtubule-associated protein 2
MDA: malondialdehyde
NACA: N-acetylcysteine amide
NCOA4: nuclear receptor coactivator 4
NRF2: nuclear factor erythroid 2-related factor 2
PBS: phosphate-buffered saline
PCR: polymerase chain reaction
PD: Parkinson’s disease
PDB: Protein Data Bank
PVDF: polyvinylidene fluoride
PUFA: polyunsaturated fatty acid
ROS: reactive oxygen species
RPA: rhodamine B-[(1,10-phenanthrolin-5-yl)aminocarbonyl] benzyl ester (RPA)
SEM: standard error of the mean
SH-SY5Y: human neuroblastoma SH-SY5Y cell line
SOD1: superoxide dismutase 1
SOD2: superoxide dismutase 2
TBARS: thiobarbituric acid reactive substances
TBA: thiobarbituric acid
TCA: trichloroacetic acid
TFR1: transferrin receptor 1
TH: tyrosine hydroxylase

## Unpublished observations and personal communications

Not Applicable

---

**Figure S1.**
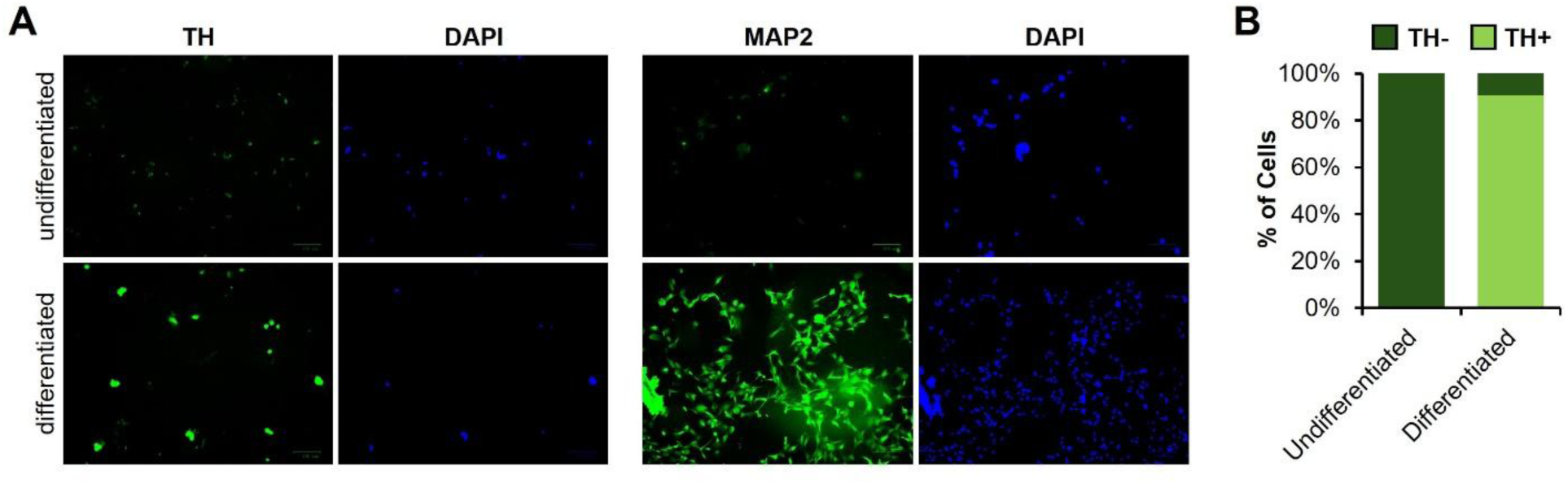
Differentiation of SH-SY5Y cells to dopaminergic neurons. **(A)** Representative immunofluorescence imaging and **(B)** quantification of neuron differentiation into dopaminergic neurons using the neuronal marker MAP2 and dopaminergic marker Tyrosine Hydroxylase (TH).

**Figure S2.**
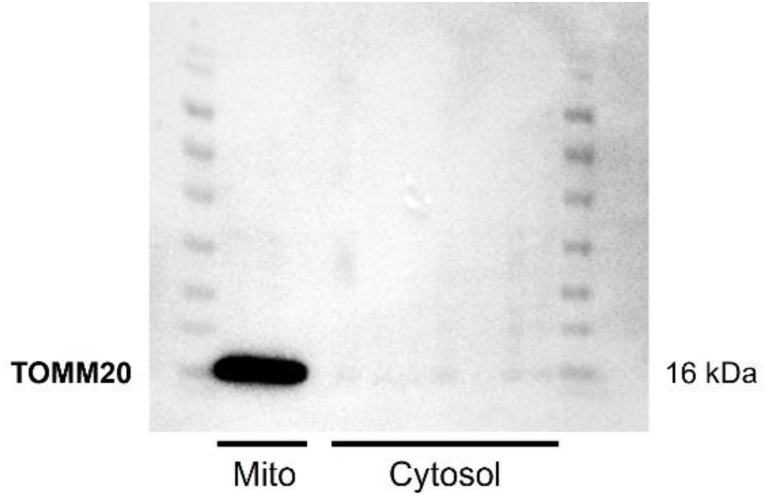
Mitochondrial isolation. Mitochondria were isolated with high purity via differential centrifugation as determined by expression of the mitochondrial-localized protein TOMM20 via western blot.

**Figure S3.**
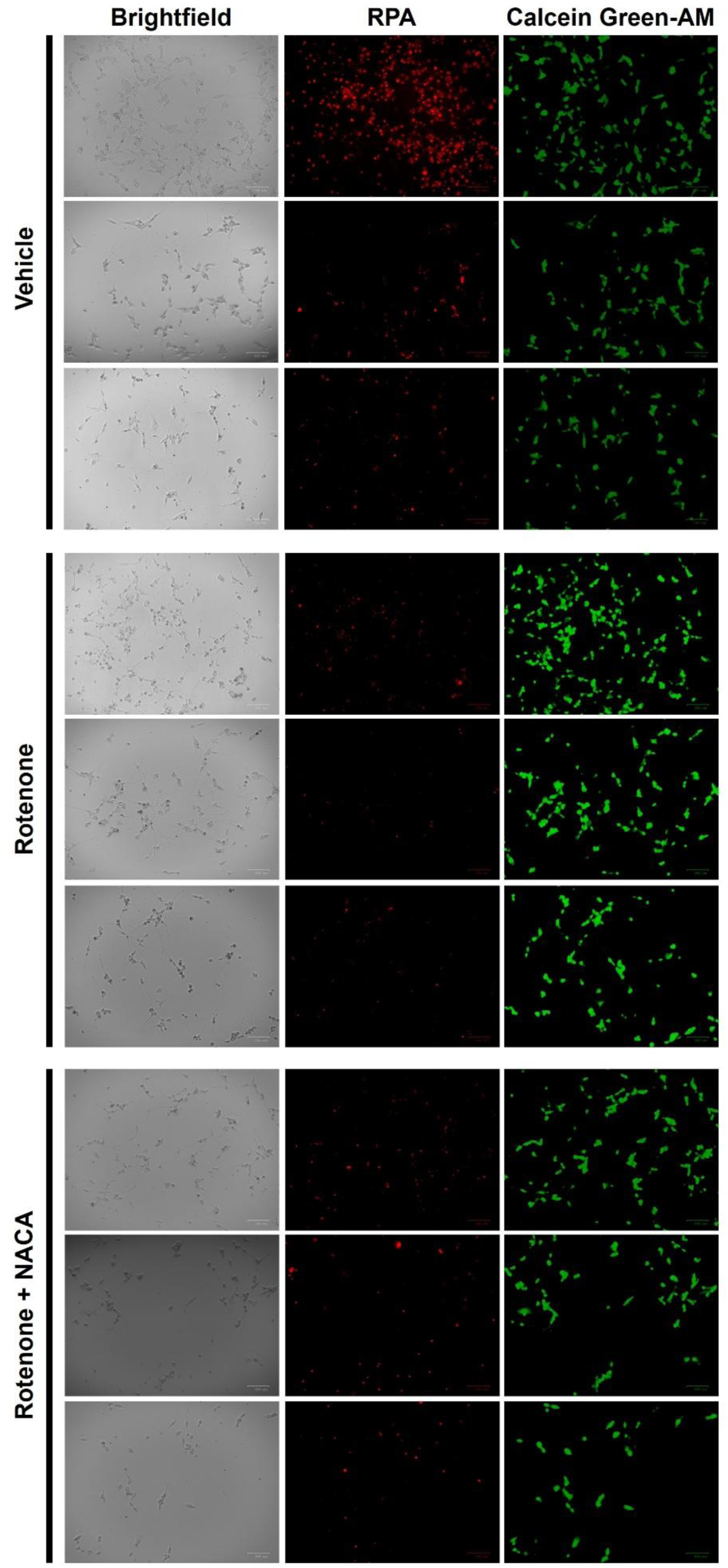
Visualization of labile iron. Labile iron in the mitochondria and cytosol was imaged in a fluorescence quenching assay using the iron-responsive fluorescent dyes RPA and calcein green-AM, respectively in SH-SY5Y cells treated with either DMSO vehicle, rotenone (25 nM), or rotenone (25 nM) in the presence of NACA (250 µM) for 24 hrs. Images were analyzed using ImageJ.

**Figure S4.**
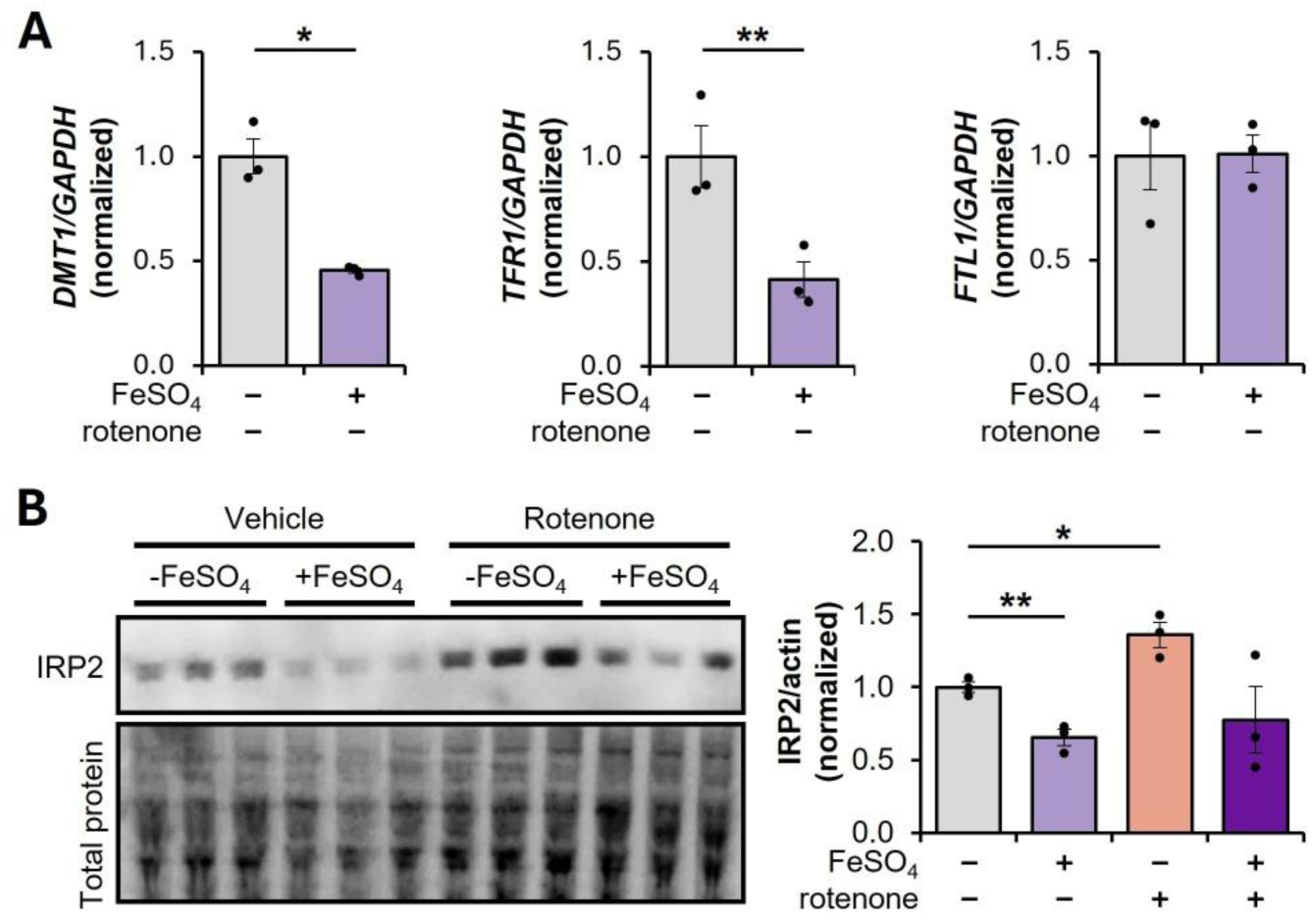
Iron overload induces cytosolic iron accumulation. **(A)** Quantification of IRE-dependent gene transcription by qPCR upon addition of iron sulfate (250 µM) for 24 hours. **(B)** Representative western blot images and densitometry (relative to actin and normalized to vehicle DMSO control) of protein expression of IRP2 in the presence or absence of rotenone (25 nM) with addition of iron sulfate (250 µM) for 24 hrs. *p < 0.05, **p < 0.01

**Figure S5.**
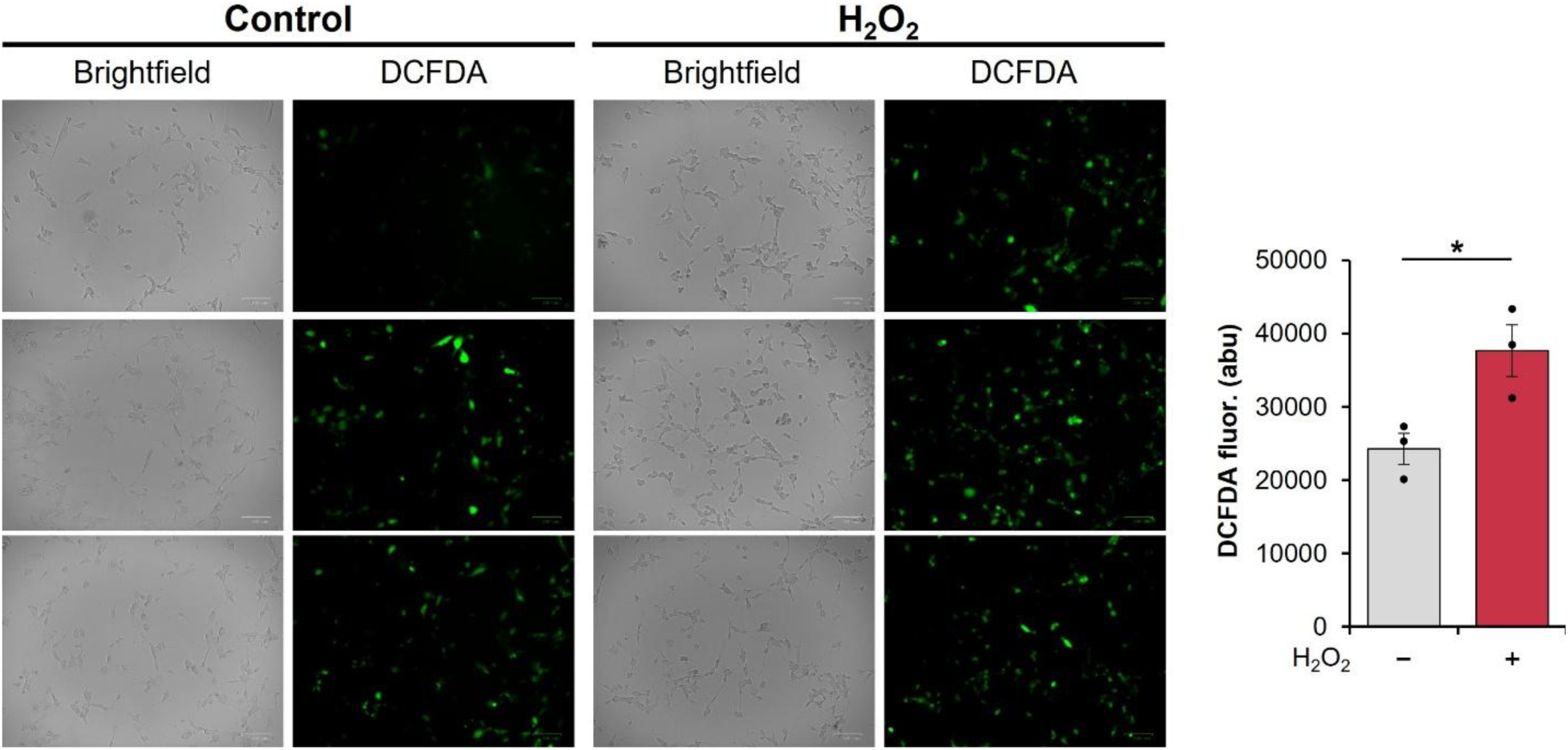
Hydrogen peroxide increases intracellular ROS. DCFDA fluorescence imaging of SH-SY5Y cells in the presence or absence of hydrogen peroxide as a positive control. Images were quantified via ImageJ. *p < 0.05

**Figure S6.**
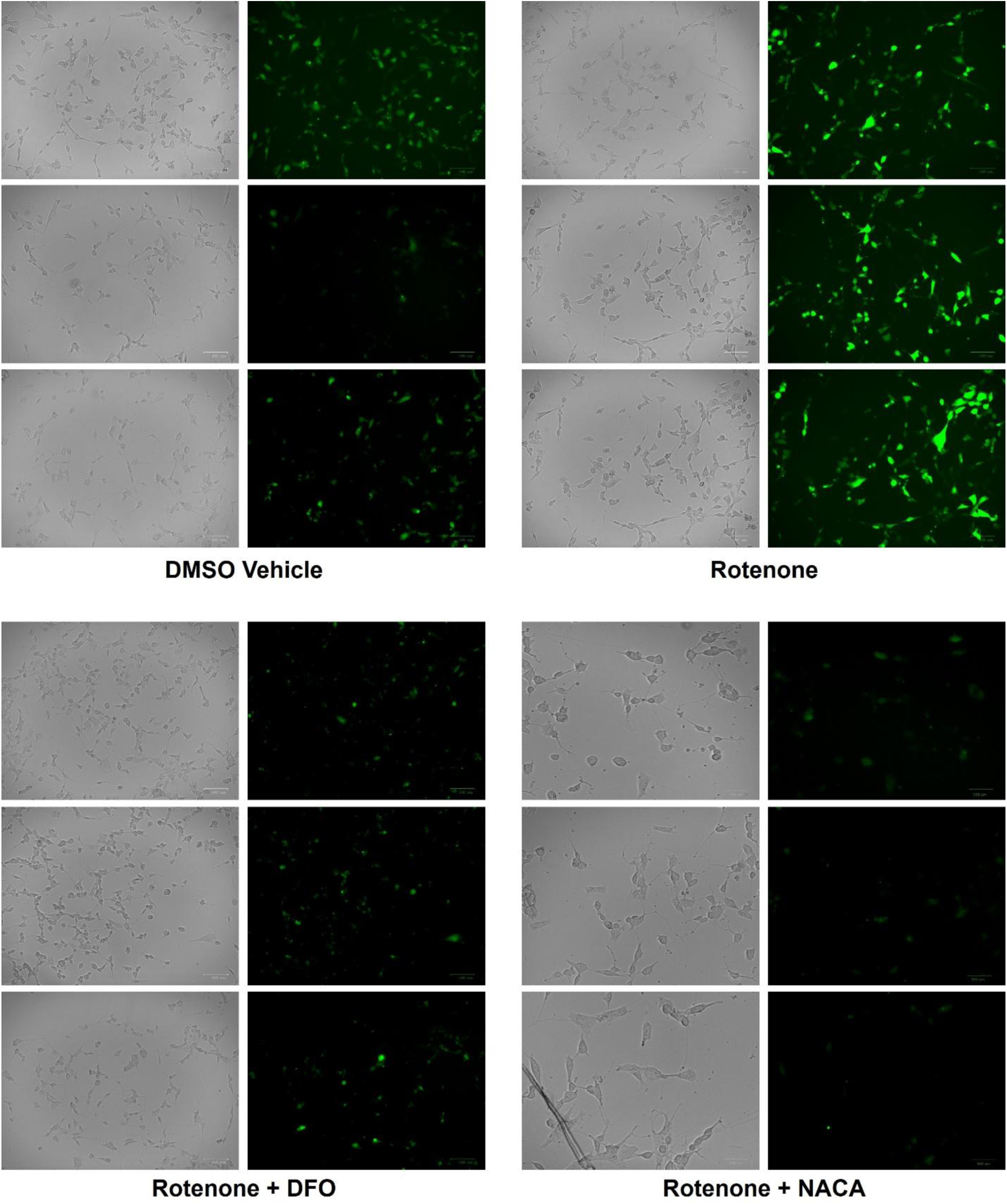
Visualization of ROS using a DCFDA Assay. DCFDA (green) was used as a readout for ROS generation in SH-SY5Y cells treated with either DMSO vehicle, rotenone (25 nM), DFO (50 µM) in the presence of rotenone (25 nM), or NACA (250 µM) in the presence of rotenone (25 nM) for 24 hrs. Brightfield images of the corresponding fluorescent images are also shown. Images were analyzed using ImageJ.

